# Cellular heterogeneity of the developing worker honey bee (*Apis mellifera*) pupa: a single cell transcriptomics analysis

**DOI:** 10.1101/2023.03.20.533557

**Authors:** Anirudh Patir, Anna Raper, Robert Fleming, Beth EP Henderson, Lee Murphy, Neil C Henderson, Emily Clark, Tom C Freeman, Mark W Barnett

**Affiliations:** The Roslin Institute, University of Edinburgh, Easter Bush, Midlothian, UK EH25 9RG; Centre for Inflammation Research, University of Edinburgh, The Queen’s Medical Research Institute, Edinburgh BioQuarter, 47 Little France Crescent, Edinburgh, UK EH16 4TJ; MRC Institute of Genetics and Cancer, University of Edinburgh, Edinburgh, UK; Edinburgh Clinical Research Facility, Welcome Trust CRF Building, University of Edinburgh, Western General Hospital, Edinburgh, UK EH4 2XU5; Beebytes Analytics CIC, The Roslin Innovation Centre, The Charnock Bradley Building, University of Edinburgh, Easter Bush, Midlothian EH25 9RG

**Keywords:** Honey bee, *Apis mellifera*, single cell RNA-Seq, network analysis, metamorphosis, development

## Abstract

It is estimated that animals pollinate 87.5% of flowering plants worldwide and that managed honey bees (*Apis mellifera*) account for 30-50% of this ecosystem service to agriculture. In addition to their important role as pollinators, honey bees are well-established insect models for studying learning and memory, behaviour, caste differentiation, epigenetic mechanisms, olfactory biology, sex determination and eusociality. Despite their importance to agriculture, knowledge of honey bee biology lags behind many other livestock species. In this study we have used scRNA-Seq to map cell types to different developmental stages of the worker honey bee (prepupa at day 11 and pupa at day 15), and sought to determine their gene signatures and thereby provide potential functional annotations for as yet poorly characterized genes. To identify cell type populations we examined the cell-to-cell network based on the similarity of the single-cells’ transcriptomic profiles. Grouping similar cells together we identified 63 different cell clusters of which 15 clusters were identifiable at both stages. To determine genes associated with specific cell populations or with a particular biological process involved in honey bee development, we used gene co-expression analysis. We combined this analysis with literature mining, the honey bee protein atlas and Gene Ontology analysis to determine cell cluster identity. Of the cell clusters identified, 9 were related to the nervous system, 7 to the fat body, 14 to the cuticle, 5 to muscle, 4 to compound eye, 2 to midgut, 2 to hemocytes and 1 to malpighian tubule/pericardial nephrocyte. To our knowledge, this is the first whole single cell atlas of honey bees at any stage of development and demonstrates the potential for further work to investigate their biology of at the cellular level.

## Introduction

The western honey bee, *Apis mellifera*, is valued for the pollination services it provides to many crops and wild flowers (Kleijn *et al.*, 2015; Breeze *et al.*, 2011; Ollerton *et al.*, 2011; Gallai *et al.*, 2009; Klein *et al.*, 2007; Corbet *et al.*, 1991) as well as *for* its production of honey and wax (Carreck, 2018; Hepburn et al., 1991). Globally there are 11 species of honey bee (Arias and Sheppard, 2005; Engel, 1999) whose distribution is restricted to Asia with the exception of the western honey bee found all over the world and indigenous to Africa, the Middle East and Europe (Seeley, 1985; Ruttner, 1988). Despite the diversity of honey bee species in Asia, the world’s beekeeping industry is based almost entirely on one species, *Apis mellifera.* In addition to their importance to agriculture and the economy, honey bees represent a useful model organism for many areas of research (Dearden *et al.*, 2009). Although less complex than mammals, honey bees possess a highly evolved social structure, a wide range of behaviours, complex communication and can learn and remember colours, shapes, fragrances and location of sources of forage (Dearden et al., 2009). Similar to the best studied model organism in the phylum Arthropoda, *Drosophila melanogaster*, honey bees are also considered a good model for understanding cognition as they possess a range of complex social and navigational behaviour with a brain that contains ~1 million neurons and are used as a model for olfactory learning (Menzel, 2012). Alternatively, as honey bees and *Drosophila* are over 300 million years diverged (Misof et al., 2014) there are many biological differences between them including eusociality, haplodiploidy, multiple discrete phenotypes from a single genotype (polyphenisms) and symbolic language (Tautz, 2008). For example, honey bees are excellent models to study polyphenism. Worker honey bees switch between different in-hive tasks eventually progressing to foraging, allowing the mechanisms required for behavioural plasticity of major life history changes to be studied (Elekonich and Roberts, 2005; Simpson et al., 2011) whilst the differential development of queen bees and worker bees is solely dependent on diet (Slater et al., 2020).

The genome for the western honey bee was first published in 2006 by the Honey Bee Genome Sequencing Consortium. This was later improved upon by Elsik et al., 2014 who found c.5,000 more protein-coding genes, 50% more than previously reported. Wallberg et al., 2019 reported a further improvement using Pac-Bio long-reads (Amel_HAv3.1). Parallel to annotating the genome, efforts have also been made to associate phenotypes with genes using omic analyses. Studies have examined changes in gene expression associated with different treatments (pheromones and pesticide) and how they relate to behaviour, phenotype and changes associated with the colony e.g. queen loss (Ma et al., 2019; Chaimanee and Pettis, 2019; Christen et al., 2016). Pheromone and pesticide treatments have also been studied in combination with various conditions [e.g. with seasonal changes (Jeon et al., 2020), infections from Varroa (Navajas et al., 2008; Zhang et al., 2010; Morfin et al., 2019) and Nosema (Badaoui et al., 2017; Li et al., 2016; Azzouz-Olden et al., 2018)]. Mechanisms underlying developmental processes such as embryogenesis, ageing and caste determination have also been analysed (Yin et al., 2018; Evans and Wheeler, 1999; He et al., 2019; Tsuchimoto et al., 2004; Azevedo et al., 2011). Whilst some of the aforementioned experiments have derived transcriptomic data from whole honey bees others have studied tissue-specific differences e.g. analysis of differences in alternate splicing patterns between the brain and fat body (Kannan et al., 2019; Wang et al., 2012; Zayed and Robinson, 2012). However, a comprehensive tissue/cell atlas of the developing honey bee is still lacking.

Bulk tissue transcriptomics atlases have been used effectively to annotate and assign function to poorly annotated genes in pig, sheep, mice, humans and *Drosophila melanogaster* (Freeman et al., 2012; Clark et al., 2017; Su et al., 2002; Chintapalli et al., 2007; Leader et al., 2018). scRNA-Seq enables the classification of cell subtypes and differentiation trajectories that is challenging with solely a bulk RNA-Seq strategy. Single-cell expression atlases have been derived from several tissues like the Tabula Muris which spans 100,000 cells across 20 mouse tissues (Tabula Muris Consortium, 2018). Other efforts like the Fly Cell Atlas have conducted exhaustive scRNA-Seq studies on individual tissues providing a comprehensive atlas, e.g. for the brain (Davie et al., 2018) and midgut (Hung et al., 2020). Studies have also tracked the development of various organisms including Drosophila (Karaiskos et al., 2017), Zebrafish (Raj et al., 2018), cnidarian (Sebe-Pedros et al., 2018) and C. elegans (Packer et al., 2019). Such studies have demonstrated the sensitivity of scRNA-Seq data in tracking cell types and their lineages while also identifying their gene signatures and how conserved they are across species. We have followed a similar approach to construct a single-cell atlas spanning two developmental stages of the worker honey bee (prepupa at day 11 and pupa at day 15). To identify cell types associated with each stage and track them through development we examined the similarity between cells based on their gene expression using a cell-to-cell network, thus revealing different cell types (which were closely connected in the network). Similarly, by using gene co-expression network (GCN) analysis, we identified coexpressed genes i.e. genes sharing a similar expression profile across samples which were likely representative of a common biology as has been shown previously (Patir et al., 2020; Patir et al., 2019). Gene signatures for the various biology associated with cell types from worker honey bee pupae across developmental stages were identified. To our knowledge this is the first analysis of the honey bee transcriptome at single cell level resolution.

## Material and Methods

### Whole *Apis mellifera* pupae cell dissociation and sorting

Honey bees are holometabolous and worker prepupae at day 11 (S1) and pupae at day 15 (S2) were chosen for this study in order to capture the key developmental stages between capping of the larval cell (day 9) and the emergence of the imago on day 21 (Oertal, 1930) (**Figure 1A**). To gather samples, a piece of brood comb containing appropriately staged pupae was collected from a single honey bee colony at the Easter Bush Campus apiary in August 2018. Pupae were removed from the comb and placed in microcentrifuge tubes on ice. Each pupa was placed in 0.5 ml HyQTase (GE Healthcare, Chicago, Illinois, USA), finely chopped with small spring scissors for one minute and incubated for 5 min at 25°C. Samples of each stage were centrifuged at 400 RCF for 5 minutes at 4°C. Cell pellets were resuspended in 1 ml WH2 medium by drawing liquid into and out of pipette tip fifteen times (Goblirsch *et al.*, 2013). Samples (n = 4 per stage) were pooled (total volume 4ml), and the cells passed into a 5ml tube through a 70 μm strainer cap (Becton, Dickinson and Company, New Jersey, USA) to remove debris and aggregated cells. Following centrifugation of the filtered cells at 400 RCF for 5 minutes at 4°C, the supernatant was discarded and the cells resuspended in 2ml WH2 medium. After further centrifugation at 400 RCF for 5 min at 4°C, cells were resuspended in 1 ml WH2 medium and stained with 1:2,000 Sytox Red (Thermo Fisher, Waltham, Massachusetts) for downstream cell viability analysis during cell sorting. Gating strategies sorted cells on the basis of their size (forwards vs side scatter area to exclude debris), single cells (forward scatter area vs height to exclude doublet cells) and viability using a 633nm laser and 660/20 band pass emission filter on an Aria IIIu FACS (Becton, Dickinson and Company, New Jersey, USA (Figure 1B and C). Before sequencing the cells were counted and tested again for viability using a TC20 automated cell counter (Bio-Rad, Hercules, California, USA).

**Figure 1.**
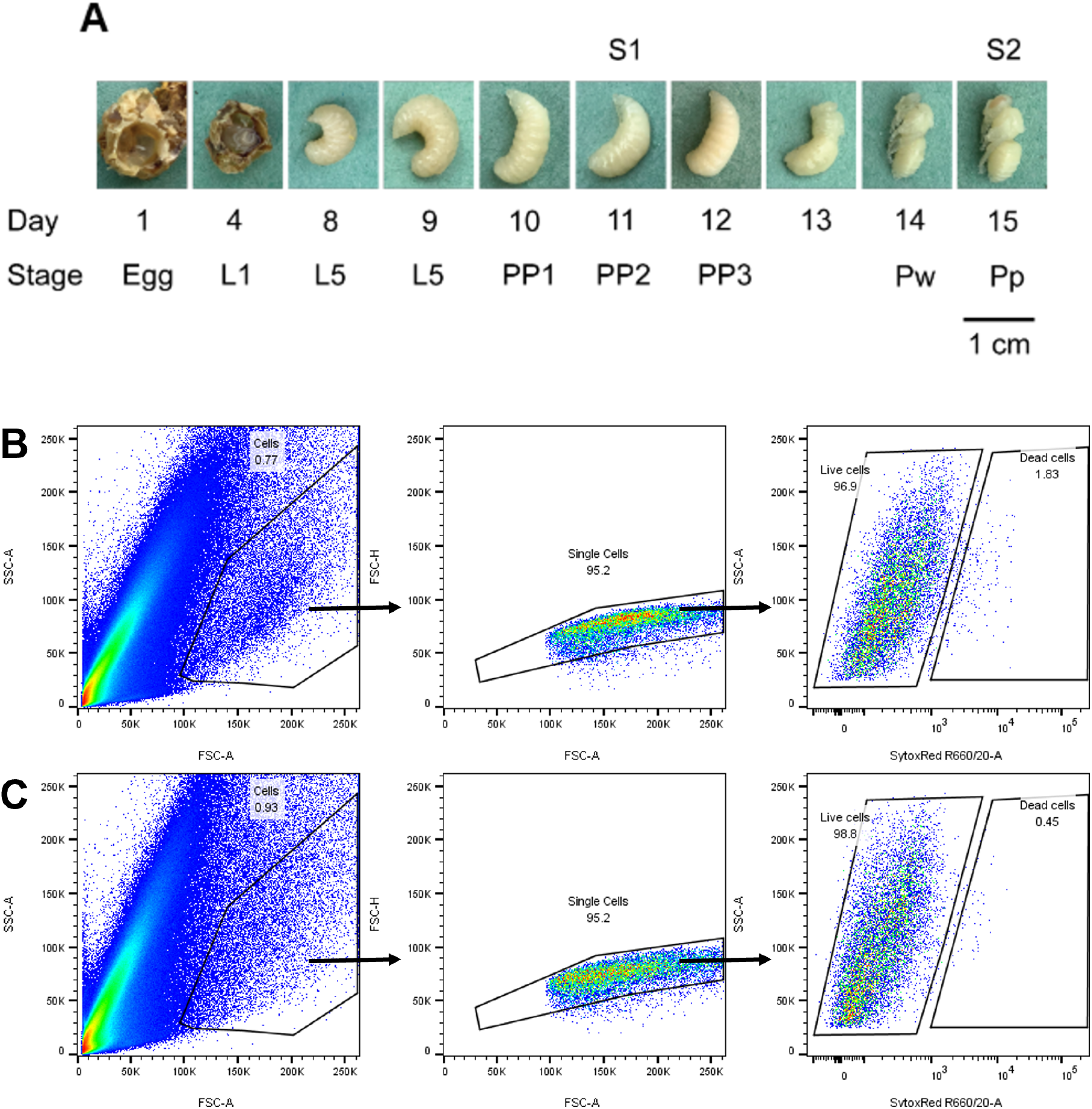
Worker honey bee development and FACS. (A) Development of honey bee worker from egg to Day 15 pupa. Queen bee was trapped on a broodless drawn broodframe in a queen excluder cage for 1 day and removed, samples of eggs, larvae and pupae were taken at one day intervals from frame within excluder cage after queen removal. L1 = 1^st^ larval instar, L5 = 5^th^ larval instar, PP1 = prepupal phase 1, PP2 = prepupal phase 2, PP3 = prepupal phase 3, Pw= white eyed pupa, Pp = pink eyed pupa. S1 and S2 were stages analysed for single cell transcriptomics. **Representative gating strategy for live single cell sort of stage 1 (B) and stage 2 (C) bee pupae**. The cells gate was defined on size and granularity and then single cells were defined using forward scatter area verses height. Live cells were then sorted by discriminating SytoxRed positive cells.

### Single-cell RNA-Seq data generation, processing and quality control

Approximately 7,000 cells at each stage were used for cDNA library preparation using the Chromium platform v2.0 (10X Genomics, Pleasanton, California, USA), as per the manufacturer’s instructions. Library quality was confirmed with a LabChip Gx24 bioanalyzer (PerkinElmer, Waltham, Massachusetts, USA). Sequencing (75bp paired-end) was performed using an Illumina NextSeq550 platform using a Mid Output 150 cycle flow cell (Clinical Research Facility, the University of Edinburgh).

Binary base call files were pre-processed using the Cell Ranger pipeline (10X Genomics). Reads were assigned to sample index tags to generate FASTQ files. Of the total 180 million reads generated, 69 million mapped to sample indices of prepupa (day 11) and 55 million to pupa (day 15). For read alignment, the recent *Apis mellifera* reference genome (Amel_HAv3.1) and annotation (GFF file) were downloaded from NCBI. To keep compatibility with Cell Ranger, the GFF file was converted to GTF using the Cufflinks software suite (Tuxedo) (Trapnell et al., 2012) and filtered for non-protein coding regions.

The resultant GTF file and reference genome were used to generate an expression matrix for each sample. Raw expression matrices were quality controlled and analysed using the Seurat package v2 in R (Stuart et al., 2019). Data from the two developmental stages were first merged. Cells having a low number of UMI reads ≤700 and ≥10% being mitochondrial were filtered out. Furthermore, genes expressed in ≤ 3 cells were removed. The data was log-normalized and genes having the most variable expression across cells were identified, i.e. possessing a standard deviation >0.5 and an average expression between 0.0125 and 3. Effects from technical factors, including variable library sizes and percent mitochondrial UMIs, were regressed out. The scaled variables were reduced to a lower feature space using principal component (PC) analysis. The most significant PCs (61 in total, *P value* < 0.05) based on JackStraw permutations (Chung and Storey, 2015) were considered and the resultant cell vs PC matrix was loaded into the network analysis tool, Graphia (Freeman et al., 2022). A correlation (Pearson similarity coefficient) matrix was then calculated between cells comparing the PC profile of each cell. Using this cell similarity matrix, a cell-to-cell network was constructed where cells (represented by a node) were connected to the 20 most similar cells by an edge, while only considering similarities beyond a Pearson cut-off threshold r ≥ 0.77. This graph was clustered using Markov clustering algorithm (Van Dongen, 2008) with an inflation value of 1.6. Cells were further filtered to remove those with an edge degree lower than three. For statistical purposes small clusters with less than 10 cells were merged into the closest cluster with the highest sum of weighted edges.

### Gene co-expression network analysis

Gene expression modules associated with biological process and cell types were identified using gene co-expression network (GCN) analysis. For conventional transcriptomics data GCNs are widely used to capture coexpressed clusters of genes associated with a shared biological function (Patir et al., 2019; Nirmal et al., 2018; Patir et al., 2020). However, due to the inherent variability within scRNA-Seq data attributed to the transcriptional heterogeneity of cells and the technical effects of dropouts (false zero expression values) we were unable to capture these coexpressing genes as they are poorly correlated. Hence, we constructed a GCN by averaging reads across cells present in each of the cell clusters described above, thus, focusing on the inter-cell type variations, i.e., the difference amongst the cell clusters, rather than the intra-cell variation. Before averaging reads, various filters were applied to reduce the effects of technical artefacts and low-level signals. First, for a given gene and cluster, cells were assigned a zero expression value if: 1) fewer than three cells within the cluster expressed that gene, 2) the maximum expression across cells was < 0.5 logged TPM and 3) < 5% of cells within the clusters expressed that gene. Moreover, to avoid the influence of outliers or spikes in expression commonly observed in RNA-Seq data we capped the maximum expression of a gene to the 95% percentile from cells of the cluster. The gene expression from the resultant filtered data was then averaged across cells for each cluster. Where a cluster consisted of cells derived from both developmental stages, they were averaged separately for each stage. In this way, the 63 cell clusters identified from the graph analysis of cells, were expanded to 81 stage differentiated cell clusters. Consequently, an expression matrix of genes vs. cell clusters was used to generate a GCN within Graphia. Only genes with a maximum expression above 0.2 average logged TPM were considered. The k-nearest neighbour (kNN) algorithm was applied where each cell was connected to the four most similar cells provided this similarity was r ≥ 0.7. Subsequently, the graph was clustered using the Louvain cluster algorithm (Blondel et al., 2008) applied with a granularity setting of 0.65.

### Functional gene annotation using *Drosophila melanogaster* homologues

Functional annotation of gene clusters from the GCN analysis was provided based on Gene Ontology (GO) enrichment analysis and literature mining. First, each protein of the bee proteome was mapped to the most similar (E score < 10^-4^) protein in *Drosophila melanogaster* (Release 6 plus ISO1 MT) based on their sequence using BLASTp (Altschul et al., 1990). The resultant nomenclature in combination with studied honey bee genes was used to functionally annotate gene clusters. Furthermore, the *Drosophila* homologues were also used for GO enrichment analysis, this was conducted for each gene cluster using the clusterProfiler package in R (Yu et al., 2012) with the genome wide annotation for *Drosophila* (org.Dm.eg.db) as the reference GO term database (Carlson et al., 2016)

## Results

### The expanding cellular diversity of the developing pupa

We developed a cell isolation protocol (Methods) from the developmental stages (S1, prepupa at day 11; S2, pupa at day 15) (**Figure 1A**) of the honey bee which provided sufficient cell numbers and viability for processing through the 10x Chromium platform v2.0. Four prepupae or pupae samples were combined for each stage. Briefly, cells from each stage were homogenised and dissociated using HyQTase enzyme, and then resuspended in WH-2 medium. The cell solution was passed through a 70 μm strainer to filter out any cell clumps and subsequently stained using Sytox red to estimate cell viability. These cells were then sorted based on their size, granularity and staining to identify viable single-cells (**Figure 1B and 1C**). Just before library preparation, the cells went through a second round of counting and variability testing to assure sufficient cells were processed for sequencing.

Raw reads from the scRNA-Seq experiment were mapped to the NCBI based *Apis mellifera* (Amel_HAv3.1) genome using Cell Ranger pipeline from 10x. 69 million reads mapped to samples from the day 11 S1 sample and 55 million reads to the day 15 S2 sample. The data was subjected to various quality control measures to remove outlier samples and genes with negligible expression. Only genes expressed in more than 3 cells were considered, leaving 9,119 genes for S1 and 9,309 genes for S2. Cells were filtered on their read content, removing cells with a low read count (< 700 per cell) and those with a high mitochondrial gene content (> 10%), leaving 2,148 cells from S1 and 2,178 cells from S2. As the two samples were from a single batch, datasets were merged and followed the standard scRNA-Seq pre-processing steps of normalistion and scaling (for mitochondrial content and library size). To cluster cells based on their gene expression profile, the 1,361 most variable genes were identified and were reduced using principal component (PC) analysis from which the 61 most significant PCs were inspected. These PCs were used to calculate Pearson pairwise similarity between cells across the merged dataset thereby generating a cell-to-cell similarity matrix. The matrix was used to construct a cell-to-cell network (**Figure 2**) where each node represented a cell and those having a Pearson correlation coefficient greater than r ≥ 0.77 were connected to one another by an edge. Furthermore, for each cell only the 20 nearest neighbours were considered and poorly connected cells, i.e. connected to < 3 other cells, were removed. These steps further helped in removing potential outlier cells that were dissimilar to the majority of cells. The final cell-to-cell graph consisted of 4,149 nodes (cells) (2,045 cells from S1 and 2,104 cells from S2) and 31,000 edges.

**Figure 2.**
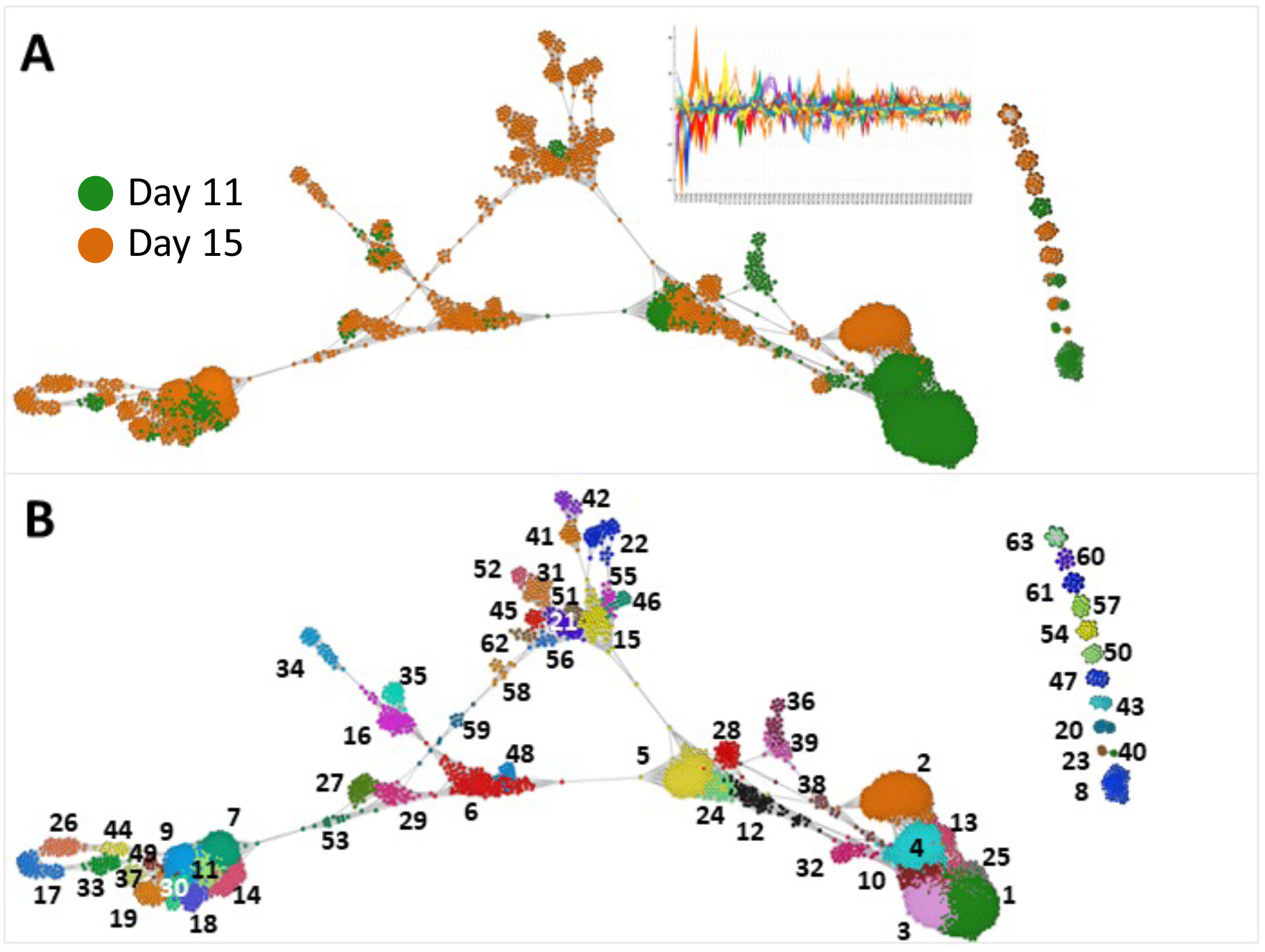
Honey bee cell populations as defined by scRNA-Seq analysis. (A) Cell-to-cell network generated by comparing the 61 most significant PCs for each cell. See insert in **(A)** showing plot of PCA profiles (y axis, each PC signified by colour) for all cells (x axis) in the graph. Each of the node represents an individual cell and the edges the 10 most significant correlations between them *r* threshold > 0.77. The graph is composed of 4,149 cells connected by 31,000 edges. In **(A)** nodes are coloured by the pupal stage from which they were derived. Note the clustering of some cells based on stage, suggesting stage-specific cell populations. In **(B),** nodes are coloured according to their cluster ID, 63 clusters being defined. The clusters disconnected from the central network are positioned on the right. Numbers indicate cluster ID.

The cell-to-cell graph consisted of one large, interconnected component and 11 smaller components. Cells from the two stages were distributed differently across the network indicative of stage-specific cell types with S2 possessing more heterogenous populations of cell types (**Figure 2A**). On studying the distribution of genes and reads across cells, cells from S2 showed a significant (1.28 times, *P value* < 10^-3^) increase in the number of genes expressed relative to S1. Clustering of the cell network resulted in 72 clusters potentially representing distinct cell types of states. To improve the statistical power of downstream analyses, smaller cell clusters with less than 10 cells were merged with a neighbouring cluster to which they were highly connected, i.e., had the highest sum total of weighted (based on the Pearson correlation) connections resulting in 63 cell clusters (**Figure 2B**). Interestingly, even though the number of cells from both stages was approximately the same, 51 clusters comprised of cells from S2, while S1 cells were only present in 30 clusters. All together, these results were indicative of the expanding cellular diversity in the developing honey bee pupa.

### Clustering of coexpressing genes and their functional annotation

To determine genes associated with specific cell populations or biological processes involved in worker honey bee development, GCN analysis was performed. The approach has been used extensively to study expression data to determine coexpressing genes, i.e., genes sharing a common expression profile across samples and which are likely to represent a common biology (Patir et al., 2019; Nirmal et al., 2020; Patir et al., 2020). Although widely used to study bulk transcriptomics data, genes have shown to be poorly correlated in scRNA-Seq data due to the complexity of single-cell biology and technical artefacts inherent to this technology (Hicks et al., 2018). Hence, we have averaged expression values across cells within a cluster to improve the stability of signals within clusters whilst also highlighting inter-cell type variation rather than the variation within a cell type (Satija and Shalek, 2014). The resultant stage-cluster vs gene expression matrix was used to calculate a gene-to-gene correlation matrix, from which we constructed a GCN. In the network, genes were connected to the four most similar genes by an edge provided they were highly correlated r ≥ 0.7. The network graph consisted of 3,994 genes which were clustered into 32 gene clusters using the Louvain clustering algorithm with a granularity of 0.65 (**Figure 3 and Table S1**). In addition to GCN analysis, differential gene expression analysis was performed using the default Wilcox test provided in Seurat to gauge the magnitude and specificity of genes towards cell-clusters based on their expression (**Table S2**)

**Figure 3.**
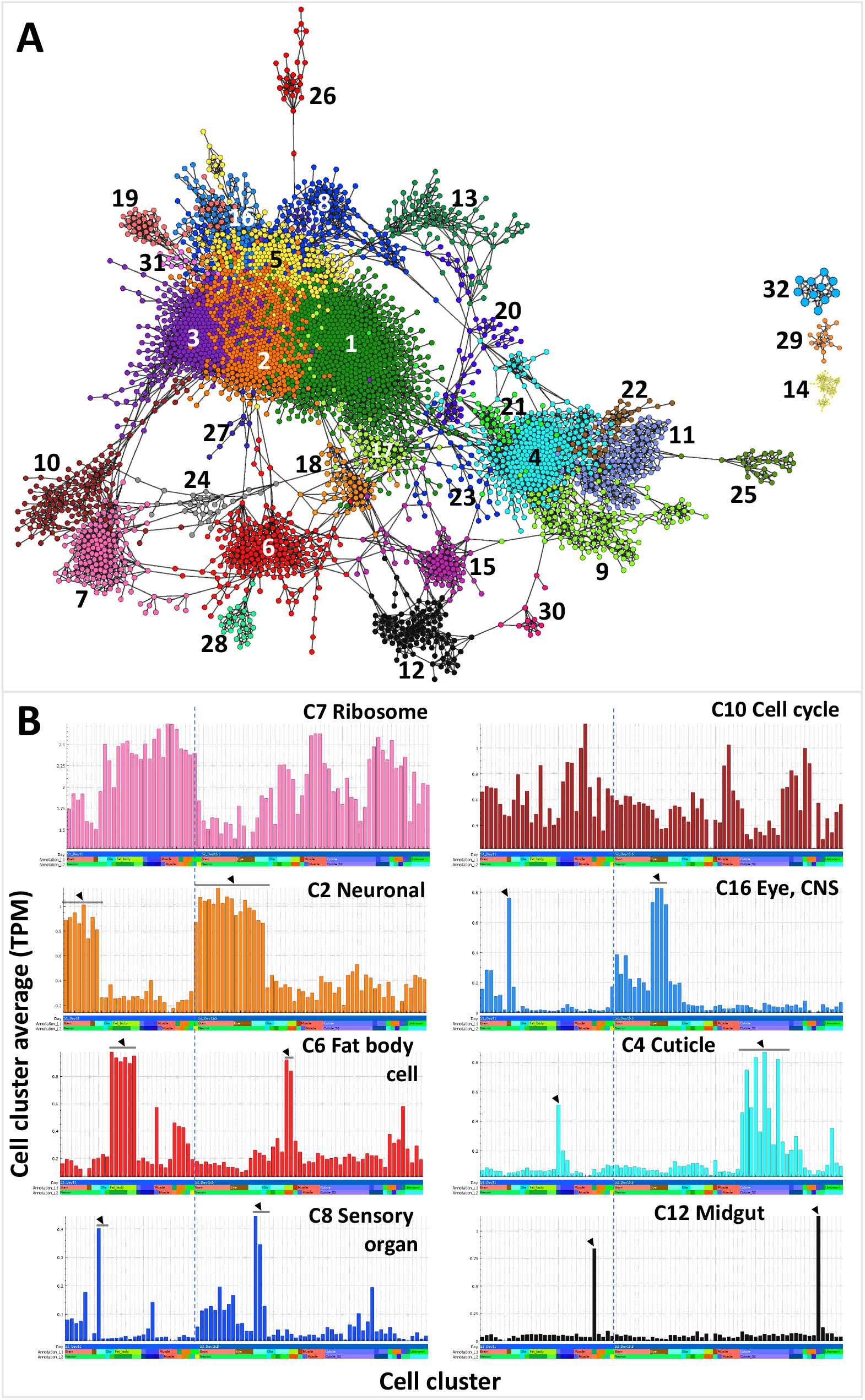
Gene correlation network analysis of expression profile of genes across cell clusters. **(A)** GCN composed of 3,994 nodes (genes) connected by 11,400 edges where *r* threshold > 0.7. Nodes are coloured according to Louvain cluster (granularity 0.65). **(B)** Average expression profile of gene clusters based on each gene’s average expression across a cluster of cells. To the left of the dotted line are cell clusters from the day 11 pre-pupa and on the right of the line are cell clusters from the day 15 pupa. Clusters of cells have been grouped based on similarity.

Tissues, cell types and biological processes corresponding to the clusters of genes in the GCN were identified from GO enrichment (**Table S1**), public resources and literature mining (**Table S3**), the final annotation of which is summarised in figure 4 & table S3. The enrichment analysis was performed on each gene cluster based on the *Drosophila melanogaster* GO reference database. For this analysis, honey bee genes based on their corresponding proteins were first mapped to the *Drosophila melanogaster* proteome using blastp (Altschul et al., 1997) where the most similar mapping was considered for a gene. 26 clusters were found to be enriched in various GO terms (*adj. P value* < 0.05) (**Table S1**). For literature mining previous publications and resources were used including the Drosophila FlyAtlas2 (Leader et al 2018) and Honey Bee Protein Atlas (Chan et al. 2013). The analyses revealed gene clusters associated with stage-dependent differences, as well as tissue/cell-specific biology, e.g. neuronal, muscle, cuticle, fat body, alimentary canal and haemolymph:

**Figure 4.**
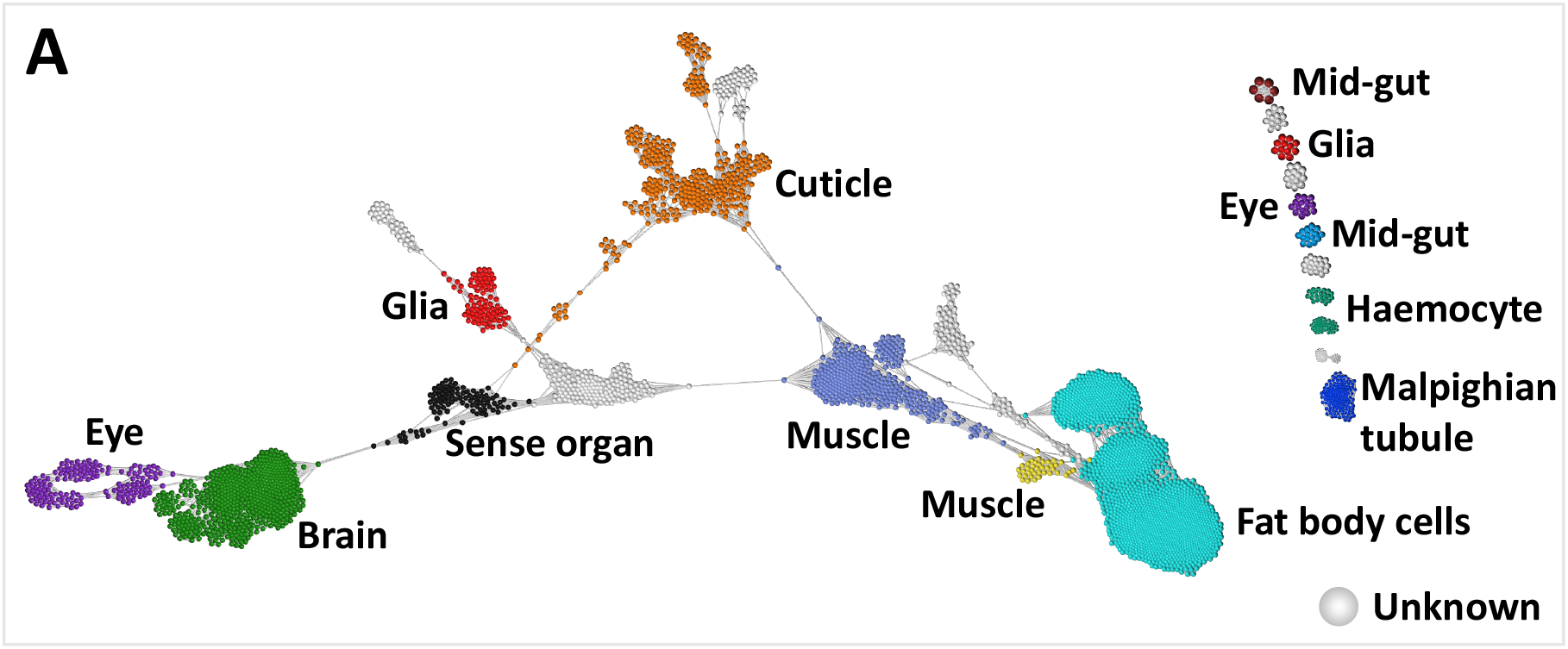
Final assignment of cell identity. The cell-to-cell network is similar to that from figure 3 where each dot represents a cell with similar cells connected to one another. However, it is overlayed with broad level annotation (colour) for the various cell clusters that have been defined based on GCN analysis, Fly Atlas2, Honey Bee Protein Atlas, and literature mining. Clusters where we could not find sufficient supporting evidence are classed as “Unknown” in grey.

#### Stage-specific clusters

The largest gene cluster, cluster 1 comprised of 708 genes (Figure 3) with a higher expression in cells from S2 relative to S1. GO terms enriched in these genes included those related to development, the top three GO terms being “post-embryonic animal morphogenesis” (*adj. P value* = 2.29×10^13^), “instar larval or pupal morphogenesis” (*adj. P value* = 3.35×10^13^) and “regulation of intracellular signal transduction” (*adj. P value* = 5.39×10^13^).

#### Neuronal related cell clusters

Three gene clusters (gene clusters 2, 3 and 5) contained genes associated with various neuronal biology and were highly expressed in 17 cell clusters identified as being related to neurons and sense organs. All of the cell clusters identified as neuronal or sense organ related expressed both synapsin and the nicotinic acetylcholine receptor alpha 1 subunit, while those annotated as only neuronal expressed the NR1 subunit of the NMDA receptor. Some genes were expressed differentially across developmental stages, including genes from the family of G protein-coupled receptors that bind octopamine and/or tyramine. Octopamine is widely distributed in the nervous system of invertebrates where it acts as a neurotransmitter (Verlinden et al., 2010) and is thought to be the functional homologue of vertebrate adrenergic transmitters. On examining the different classes of these G protein-coupled receptors (Sinakevitch et al., 2017) in invertebrates, OA1 receptor showed a high expression in cell clusters 37 and 60 containing S2 cells, AmTAR1 was highly expressed in cell cluster 33 having cells from both pupal stages, whilst AmTARII showed a high expression in cell clusters 9 and 11 of S1 stage

#### Glial related cell clusters

Glial cell*s* have an essential role in the development of neurons and are involved in regulation of synaptic plasticity, provide trophic support to neurons and contribute to the blood-brain barrier (Shah et al., 2018). In the honey bee these cells can be labelled using a serum raised against the *Drosophila* glial transcription factor *repo* (Shah et al., 2018), *repo* was highly expressed in several non-neuronal cell clusters (6, 16, 34, 35, 48 and 61) identifying them as potentially representing glia or glial-related cells. Gene cluster 16 was found to be associated with these cell clusters. Further subclassification of these cells was revealed through genes linked with astrocytes in *Drosophila*, like *Eaat2* and GABA transporters (Gat-a and Gat-1b) highly expressed in cell cluster 35 (Freeman, 2015).

#### Sensory organ and compound eye related cell clusters

A higher average expression of genes from gene cluster 8 was observed in sensory organs relative to neuron related cell clusters. GO terms enriched in genes from this cluster were associated with ciliary biology, the most significant terms being “cilium organization” (*adj. P value* = 6.48×10^24^), “cilium assembly” (*adj. P value* = 8.53×10^24^) and “plasma membrane bounded cell projection assembly” (*adj. P value* = 4.36×10^21^). The modified primary cilium is a structure common to all peripheral sensory neurons in arthropods with the exception of photoreceptors (Keil, 2012), suggesting that cell clusters 27, 29 and 53 were related to sense organs other than the compound eye and ocelli. Four cell clusters (clusters 26, 33, 44 and 49) identified as neural were associated with the compound eye. Genes from gene cluster 16 (80 genes) were specifically expressed in these eye related cell clusters with genes associated with this tissue e.g. *AmPNR-like* (LOC413558) shown by *in situ* hybridization to be expressed in the developing eyes of pupae in either the photoreceptor cells or support cells (Velarde et al., 2006). LOC408804 (1-phoshatidylinositol 4,5 bisphosphate phosphodiesterase epsilon-1) was expressed in these cell clusters and in Drosophila it’s homologue (*Plc21c*) has a role in Pigment Dispersing Factor neurons in the circadian photoresponse (Ni et al., 2017). Phosrestin 2 was specifically expressed in cell clusters 17, 26 and 44, and has been associated with the visual system in honey bees where it has a role in circadian rhythms (Rodriguez-Zas et al., 2012).

#### Cuticle related cell clusters

Gene clusters 4, 9, 11, 22 and 30 included genes expressed in 20 cell clusters associated with the cuticle. Only four of these cell clusters were associated with the S1 prepupal cuticle (cell clusters 36, 38, 39, 51). This could indicate that cell populations from the S1 stage cuticle are less diverse than those from S2 which might consist of heterogenous populations of cells differentiating in different regions of the developing honey bee exoskeleton. The cuticle associated gene clusters included key enzymes in the chitin biosynthetic pathway linked to cuticle development and the moulting process e.g. LOC412215 (homologue of Drosophila gene *kkv*, a chitin synthase that catalyses the conversion of UDP-*N*-acetylglucosamine to chitin), LOC552276 (homologue of Drosophila gene *cda5*), a chitin deacetylase that catalyses the conversion of chitin to chitosan (a polymer of *β*-1,4-linked d-glucosamine residues) (Sobala and Adler, 2016) and LOC551964 (homologue of *Drosophila* gene *mmy*, an enzyme required for glycan and chitin synthesis) (Araujo et al., 2005). Chitin (the polymer of N-acetyl glucosamine) is a key component of the honey bee inner procuticle, which together with the outer epicuticle forms the exoskeleton (Locke and Krishnan, 1971) and the difference in cuticle structure in arthropods is due to the different expression of proteins (Magkrioti at al., 2004). In addition to chitin, the cuticle consists of various cuticle structural proteins some of which were present in the cuticle related gene clusters including LOC726451 (homologue of *Drosophila* gene *Cpr57A*) and *Apd-3* (Falcon et al., 2019).

#### Fat body related cell clusters

In insects this tissue has a similar role to the liver and adipose tissue of mammals as it functions as a store for excess nutrients, synthesizes most of the haemolymph proteins and is responsible for detoxification processes (Arrese and Soulages, 2010). Various genes associated with the fat body were found in gene cluster 6 (199 genes) which had a high expression in seven cell clusters. The majority of these clusters comprised cells from the S1 stage (six clusters). The gene *ilp-2* known to be expressed in both oenocytes and trophocytes (cell types found in the fat body) in the adult honey bee and was expressed in all seven fat body related cell clusters (Nilsen et al., 2011). A similar expression was observed for *mmp2* shown to be necessary and sufficient for fat body remodelling during early metamorphosis in *Drosophila* (Bond et al., 2011) and Vitellogenin receptor shown to be expressed in the fat body, ovary and head of adult worker bees (Guidugli-Lazzarini et al., 2008).

#### Haemolymph related cell clusters

In insects, haemocytes are derived from anterior mesoderm, form part of the immune system and comprise lamellocytes, crystal cells, plasmatocytes and granulocytes (Richardson et al, 2018). Granulocytes are the major phagocytic cells (Richardson et al, 2018). Gene clusters 28 and 32 contained known haemocyte markers (*hml* and *lz*) and the average expression of these genes was higher in cell clusters 20 and 43. The marker *hml* (hemolectin/hemocytin) is specifically expressed in haemocytes in *Drosophila* in embryos and larvae, while *lz* is required for the differentiation of crystal cells (Lebestky et al., 2003) and the absence of its expression results in the differentiation of a plasmatocyte. Whilst both gene were expressed highly in the haemocyte cell clusters, *lz* showed higher levels of expression in cell cluster 43, suggesting that it represented crystal cells.

#### Muscle related cell clusters

In Drosophila, somatic muscle, visceral muscle and cardiac muscle develop from the mesoderm (Gunage et al., 2017). The largest somatic muscles in the honey bee are two pairs of indirect flight muscles (dorsumventral and anterior-posterior) in the thorax that are responsible for moving the wings up and down (Snodgrass, 1910). Cell clusters 5, 12, 24, 28 and 32 were annotated as differentiating muscle cells based on expression of twist (Gunage et al., 2017), mef2 (Crittenden et al., 2018), nautilus (Abmayr and Keller, 1998), TpnT (Domingo et al., 1998), TpNI (Herranz et al., 2005), TpnCIIb (Herranz et al., 2005), myosin heavy chain (LOC409843) and myosin light chain (LOC409881) (11390828). *nautilus* is a candidate for the equivalent of the vertebrate myogenic regulatory factors (*myoD* and *Myf5*) that act as master control genes in mesoderm to initiate the first steps of somatic muscle development (Abmayr and Keller, 1998; Zammit, 2017). Expression of *nautilus* specifically in cell clusters 12, 24 and 32 indicated that these cell clusters comprised of cells differentiating into somatic muscle. Expression of twist in Drosophila is required earlier in development in mesoderm definition for specification of all muscle types, *twist* was expressed specifically in cell clusters 5, 12, 24, 28 and 32, and was also expressed in cell cluster 60 (unknown identity). Cell clusters 5, 12, 24, 28 and 32 were also associated with differentiating muscle cells based on GO analysis of gene cluster 14 whose gene showed high expression in these cells relative to other cell clusters. The top GO terms for the gene cluster 14 included “striated muscle cell differentiation” (*adj. P value* = 2.02×10^20^), “muscle structure development” (*adj. P value* = 4.12×10^20^), and “muscle cell differentiation” (*adj. P value* = 8.61×10^20^).

##### Alimentary canal

the tissue comprises four major compartments, the foregut, midgut, malpighian tubules and hind gut (Snodgrass, 1910). Genes from gene cluster 12 were highly expressed in cell clusters 8 and 50. The associated gene cluster included alpha-glucosidase I and II shown to be expressed in honey bee ventriculus (Kubota et al., 2004), as well as organic anion transporting polypeptide genes *Oatp33Ea* and *Oatp58Dc* both of which are specific to the *Drosophila* midgut of larva and adult based on the FlyAtlas 2 tissue RNA-Seq database (Leader et al., 2018). Cell cluster 63 had a high expression of genes from gene cluster 18 thought to related to malpighian tubules or pericardial nephrocytes, including *Cubilin* and *Amnionless* which in *Drosophila* mediate protein reabsorption in both malpighian tubules and pericardial nephrocytes (Zhang et al., 2013).

## Discussion

The aim of this study was to generate single cell transcriptomics data for two stages of worker honey bee development. We have generated scRNA-Seq data from a prepupal stage (day 11) and the pupal stage (day 15), these stages being selected to capture cellular diversity immediately before and after the rearrangement of the larval to adult body plan. In holometabolous insects, the larvae and adults have very different body plans enabling them to exploit different resources. Although the larvae of social insects and solitary bees have subsequently evolved to be relatively immobile, this remarkable evolutionary development contributed to holometabolous insects comprising over half of global eukaryotic diversity (Belles, 2017). Despite the importance of metamorphosis in the evolutionary success of insects, the mechanisms governing it are not completely understood. Most is known about regulation by the endocrine system which involves the hormones 20E (20-hydroxyecdysone) and JH (juvenile hormone) (Truman and Riddiford, 2019). Generally, 20E promotes moulting whilst JH inhibits metamorphosis and thus if 20E acts together with JH, moulting results in a juvenile stage and if it acts without JH, it results in metamorphosis (Truman and Riddiford, 2019). Activation of the 20E receptor complex results in the up regulation of transcription factors e.g., *HR3, HR4, HR39, Broad complex, E75* and *FTZ-F1* (King-Jones and Thummel, 2005; Nagakawa and Henrich, 2009). Activation of the JH receptor complex results in induction of *Kr-h1*, Kr-h1 subsequently represses transcription of *E93* (Belles and Santos, 2014; Urena et al., 2014). If E93 protein levels increase, metamorphosis is triggered. During metamorphosis, tissues can degenerate if they are not present in the adult (e.g. head gland), be remodelled without complete cell replacement (e.g. fat body) or generate a new adult structure (e.g. antenna, eyes, legs and wings develop from undifferentiated cells in imaginal discs) (Tettamanti and Casartelli, 2019).

Two strategies have previously been adopted by other researchers in studying development using scRNA-Seq. The first involves scRNA-Seq of whole-organisms and the second, of focussing on individual tissues. Here, we adopt the former approach which has proven useful in the exploration of cell types of model organisms of a similar scale and biological complexity, such as Cnidaria (Sebe-Pedros et al., 2018), *C. elegans* (Packer et al., 2019), and zebrafish (Farnsworth et al., 2020), where the cell diversity is largely unknown. Similar to these studies we have identified the cellular diversity across different lineages and their contribution to each pupal stage. For this study we developed a protocol that can be used to prepare single cells of honey bee worker pupae for scRNA-Seq. Further research could address a wider developmental series and ascertain the efficacy of the protocol as the cuticle toughens in the later pupal stages. It seems unlikely that the protocol would be suitable for a whole adult honey bee due to the presence of a fully developed exoskeleton, however this was not attempted by the authors. The protocol should however be effective for analysis of single cells from a dissected adult brain or other dissected tissues.

The cell-to-cell network grouped cells into 63 clusters across which cells from the two stages were differently distributed. Hence, clustering of cells revealed stage-dependent cell types/subtypes i.e. certain cell types were entirely represented by cells from a single stage while other clusters comprised cells from both stages. The majority of cell clusters were entirely comprised of cells from S2, furthermore these cells had a greater number of genes expressed relative to cells from S1. These results suggest an expanding heterogeneity for the types of cells and genes, which define them and reflect the fact that most of the organs of the adult honey bee are present at S2 whilst at S1, a lower number of larval tissues are about to be degenerated, remodelled or replaced. To study the genes that were associated with the cell clusters we developed a novel approach to improve the biological signal representing inter-cell cluster variation. Briefly, this was done by averaging the reads across cells from the same cluster and applying filters on the expression values to address certain technical artefacts within the data including spikes in expression and the variation of lowly expressed genes. The approach enabled the construction of a GCN from scRNA-Seq data, which captures inter-cell type variation while minimising intra-cell type and technical variations. The GCN comprised 32 clusters of co-expressing genes that were associated with a wide range of biology as determined using a combination of GO enrichment and literature mining to identify cell types and tissue-specific biology. Cell types and tissues identified were related to the brain, sensory organs, cuticle, muscle, fat body, blood and alimentary canal. Gene coexpression signatures were identified that were not only unique to cell clusters but also those that were shared across clusters e.g. stage and lineage specific signatures. Some cell clusters would have proved impossible to identify based on using literature for *Apis* only due to the limitations of the available resources as such it was necessary to compare to *Drosophila* where organs are evolutionary conserved and where a database for GO terms are present.

Many honey bee tissues were either not detected or not identified in our analysis e.g., endocrine system, salivary glands, hypopharyngeal glands, oesophagus, honey sac, small intestine, heart, rectum, sting, ovary. This might be because there is insufficient scientific literature relevant to these pupal stages for identification (12 of the 63 clusters remain unidentified) or it might be that the protocol was either too harsh or too gentle to obtain particular cell types. It is surprising that there are noticeably few cells from the ventriculus (mid-gut) despite the relatively large size of this organ in the adult bee, and it therefore seems likely that a harsher or longer digestion might yield more cells from the mid-gut

With the lack of a gene expression atlas for the honey bee, this study provides an initial step in determining the cellular heterogeneity, which can only be improved upon by sequencing more samples/cells, cross species comparisons and analysis of gene expression experiments. This study will be of benefit to the construction of more comprehensive gene expression atlases by demonstrating that pupae can be analysed at the single-cell level, which can be potentially extended to larvae and dissected adult organs e.g. brain. Furthermore, the dataset could be used in conducting cross-species comparisons for development, as has been done for Cnidaria (Sebe-Pedros et al., 2018), to study the evolution of certain cell types.

## Conclusions

In summary, we have demonstrated that a gene expression atlas of the whole honey bee at the level of single cells is possible at prepupal and pupal stages. We have developed approaches from single cell isolation to the analysis of the resultant scRNA-Seq data using GCN. Through this process we have identified several potential cell types and their associated gene signatures which are supported by enrichment analysis, and previous experimental evidence from the literature or databases. The gene lists associated with the cell clusters will be of benefit to future analyses, particularly for transcriptomic studies in whole pupae. Despite the global importance of bees to agriculture, this is the first whole organism RNA expression atlas in Hymenoptera. As a result, the study provides, improved knowledge of transcriptional profiles of many cell types of the worker honey bee at the pupal stage, and functional annotation of its genome.

## Author Contributions

**AP** performed cell preparations from bee pupae, bioinformatics, transcriptomics analysis to assign cell identities, assisted with beekeeping and wrote the manuscript, **AR** and **RF** assisted with experimental design and performed FACS, **BH** and **NH** prepared 10X Genomics libraries and provided advice on experimental strategy, **LM** performed short read Illumina sequencing, **EC** provided advice on manuscript preparation and helped to draft the manuscript, **TF** conceived the idea for the study and managed the project, **MB** performed cell preparations from bee pupae, transcriptomics analysis to assign cell identities, managed beekeeping and wrote the manuscript. All authors read and approved the final version of the manuscript.

## Funding

This work was funded by the Biotechnology and Biological Sciences Research Council (BBSRC) Institute Strategic Programme grant “Prediction of genes and regulatory elements in farm animal genomes” (BBS/E/D/10002070) awarded to the Roslin Institute. EC and links with EMBL-EBI were supported by BBSRC grants “Ensembl—adding value to animal genomes through high-quality annotation” (BB/S02008X/1) and “Ensembl in a new era - deep genome annotation of domesticated animal species and breeds” (BB/W018772/). N.C.H. is supported by a Wellcome Trust Senior Research Fellowship in Clinical Science (ref. 219542/Z/19/Z). The Edinburgh Clinical Research Facility is funded by the Wellcome Trust.

## Data Availability

The dataset described in this manuscript has been deposited in the National Center for Biotechnology Information BioProject database (BioProject ID: PRJEB45881).

## Conflicts of Interest

The authors declare that the research was conducted in the absence of any commercial or financial relationships that could be construed as a potential conflict of interest.

## Notes

### Competing Interest Statement

The authors have declared no competing interest.

### Summary of Updates

This version of the manuscript has been revised to correct formatting of the author affiliations (affiliations 4 & 5 had merged), and to move all the panels of figure 1 together on one page. Apologies for missing these formatting issues in the previously submitted version. The supplemental material has also been uploaded with this version.

## References

Abmayr, S.M. and Keller, C.A. 1998. Drosophila myogenesis and insights into the role of nautilus. Current Topics in Developmental Biology, 38 pp. 35–80.

Altschul, S.F., Madden, T.L., Schaffer, A.A., Zhang, J., Zhang, Z., Miller, W. and Lipman, D.J. 1997. Gapped BLAST and PSI-BLAST: A new generation of protein database search programs. Nucleic Acids Research, 25 (17), pp. 3389–3402.

Altschul, S.F., Gish, W., Miller, W., Myers, E.W. and Lipman, D.J. 1990. Basic local alignment search tool. Journal of Molecular Biology, 215 (3), pp. 403–410.

Araujo, S.J., Aslam, H., Tear, G. and Casanova, J. 2005. Mummy/cystic encodes an enzyme required for chitin and glycan synthesis, involved in trachea, embryonic cuticle and CNS development--analysis of its role in drosophila tracheal morphogenesis. Developmental Biology, 288 (1), pp. 179–193.

Arias, M.C. and Sheppard, W.S. 2005. Phylogenetic relationships of honey bees (hymenoptera:Apinae:Apini) inferred from nuclear and mitochondrial DNA sequence data. Molecular Phylogenetics and Evolution; Mol Phylogenet Evol, 37 (1), pp. 25–35.

Arrese, E.L. and Soulages, J.L. 2010. Insect fat body: Energy, metabolism, and regulation. Annual Review of Entomology, 55 pp. 207–225.

Azevedo, S.V., Caranton, O.A., de Oliveira, T.L. and Hartfelder, K. 2011. Differential expression of hypoxia pathway genes in honey bee (apis mellifera L.) caste development. Journal of Insect Physiology, 57 (1), pp. 38–45.

Azzouz-Olden, F., Hunt, A. and DeGrandi-Hoffman, G. 2018. Transcriptional response of honey bee (apis mellifera) to differential nutritional status and nosema infection. BMC Genomics, 19 (1), pp. 628–0.

Badaoui, B., Fougeroux, A., Petit, F., Anselmo, A., Gorni, C., Cucurachi, M., Cersini, A., Granato, A., Cardeti, G., Formato, G., Mutinelli, F., Giuffra, E., Williams, J.L. and Botti, S. 2017. RNA-sequence analysis of gene expression from honeybees (apis mellifera) infected with nosema ceranae. PloS One, 12 (3) pp. e0173438.

Belles, X. 2017. MicroRNAs and the evolution of insect metamorphosis. Annual Review of Entomology, 62 pp. 111–125.

Belles, X. and Santos, C.G. 2014. The MEKRE93 (methoprene tolerant-kruppel homolog 1-E93) pathway in the regulation of insect metamorphosis, and the homology of the pupal stage. Insect Biochemistry and Molecular Biology, 52 pp. 60–68.

Blondel, V.D., Guillaume, J., Lambiotte, R. and Lefebvre, E. 2008. Fast unfolding of communities in large networks. Journal of Statistical Mechanics: Theory and Experiment, (October) pp. P10008.

Bond, N.D., Nelliot, A., Bernardo, M.K., Ayerh, M.A., Gorski, K.A., Hoshizaki, D.K. and Woodard, C.T. 2011. ssFTZ-F1 and matrix metalloproteinase 2 are required for fat-body remodeling in drosophila. Developmental Biology, 360 (2), pp. 286–296.

Breeze, T.D., Bailey, A.P., Balcombe, K.G. and Potts, S.G. 2011. Pollination services in the UK: How important are honeybees? Agriculture, Ecosystems & Environment, 142 (3), pp. 137–143.

Carlson, M.R.J., Pagès, H., Arora, S., Obenchain, V. and Morgan, M. 2016. Genomic annotation resources in R/bioconductor. In: Ewy Mathé and Sean Davis. ed. Statistical Genomics: Methods and Protocols New York, NY:Springer New York.

Carreck, N.L. 2018. Special issue: Honey. Journal of Apicultural Research, 57 pp. 1–4.

Chaimanee, V. and Pettis, J.S. 2019. Gene expression, sperm viability, and queen (apis mellifera) loss following pesticide exposure under laboratory and field conditions. Apidologie, 50 pp. 304–316.

Chan, Q.W., Chan, M.Y., Logan, M., Fang, Y., Higo, H. and Foster, L.J. 2013. Honey bee protein atlas at organ-level resolution. Genome Research, 23 (11), pp. 1951–1960.

Chintapalli, V.R., Wang, J. and Dow, J.A. 2007. Using FlyAtlas to identify better drosophila melanogaster models of human disease. Nature Genetics, 39 (6), pp. 715–720.

Christen, V., Mittner, F. and Fent, K. 2016. Molecular effects of neonicotinoids in honey bees (apis mellifera). Environmental Science & Technology, 50 (7), pp. 4071–4081.

Chung, N.C. and Storey, J.D. 2015. Statistical significance of variables driving systematic variation in high-dimensional data. Bioinformatics (Oxford, England), 31 (4), pp. 545–554.

Clark, E.L., Bush, S.J., McCulloch, M.E.B., Farquhar, I.L., Young, R., Lefevre, L., Pridans, C., Tsang, H.G., Wu, C., Afrasiabi, C., Watson, M., Whitelaw, C.B., Freeman, T.C., Summers, K.M., Archibald, A.L. and Hume, D.A. 2017. A high resolution atlas of gene expression in the domestic sheep (ovis aries). PLoS Genetics, 13 (9) pp. e1006997.

Corbet, S.A. 1991. Bees and the pollination of crops and wild flowers in the european community. Bee World, 72 pp. 47–59.

Crittenden, J.R., Skoulakis, E.M.C., Goldstein, E.S. and Davis, R.L. 2018. Drosophila mef2 is essential for normal mushroom body and wing development. Biology Open, 7 (9), pp. 10.1242/bio.035618.

Davie, K., Janssens, J., Koldere, D., De Waegeneer, M., Pech, U., Kreft, L., Aibar, S., Makhzami, S., Christiaens, V., Bravo Gonzalez-Blas, C., Poovathingal, S., Hulselmans, G., Spanier, K.I., Moerman, T., Vanspauwen, B., Geurs, S., Voet, T., Lammertyn, J., Thienpont, B., Liu, S., Konstantinides, N., Fiers, M., Verstreken, P. and Aerts, S. 2018. A single-cell transcriptome atlas of the aging drosophila brain. Cell, 174 (4), pp. 982–998.e20.

Dearden, P.K., Duncan, E.J. and Wilson, M.J. 2009. The honeybee apis mellifera. Cold Spring Harbor Protocols, 2009 (6), pp. pdb.emo123.

Domingo, A., Gonzalez-Jurado, J., Maroto, M., Diaz, C., Vinos, J., Carrasco, C., Cervera, M. and Marco, R. 1998. Troponin-T is a calcium-binding protein in insect muscle: In vivo phosphorylation, muscle-specific isoforms and developmental profile in drosophila melanogaster. Journal of Muscle Research and Cell Motility, 19 (4), pp. 393–403.

Elekonich, M.M. and Roberts, S.P. 2005. Honey bees as a model for understanding mechanisms of life history transitions. Comparative Biochemistry and Physiology.Part A, Molecular & Integrative Physiology, 141 (4), pp. 362–371.

Elsik, C.G., Worley, K.C., Bennett, A.K., Beye, M., Camara, F., Childers, C.P., de Graaf, D.C., Debyser, G., Deng, J., Devreese, B., Elhaik, E., Evans, J.D., Foster, L.J., Graur, D., Guigo, R., HGSC production teams, Hoff, K.J., Holder, M.E., Hudson, M.E., Hunt, G.J., Jiang, H., Joshi, V., Khetani, R.S., Kosarev, P., Kovar, C.L., Ma, J., Maleszka, R., Moritz, R.F., Munoz-Torres, M.C., Murphy, T.D., Muzny, D.M., Newsham, I.F., Reese, J.T., Robertson, H.M., Robinson, G.E., Rueppell, O., Solovyev, V., Stanke, M., Stolle, E., Tsuruda, J.M., Vaerenbergh, M.V., Waterhouse, R.M., Weaver, D.B., Whitfield, C.W., Wu, Y., Zdobnov, E.M., Zhang, L., Zhu, D., Gibbs, R.A. and Honey Bee Genome Sequencing Consortium. 2014. Finding the missing honey bee genes: Lessons learned from a genome upgrade. BMC Genomics, 15 pp. 86–86.

Engel, M.S. 1999. The taxonomy of recent and fossil honey bees (hymenoptera: Apidae: Apis). Journal of Hymenoptera Research, 8 pp. 165–196.

Evans, J.D. and Wheeler, D.E. 1999. Differential gene expression between developing queens and workers in the honey bee, apis mellifera. Proceedings of the National Academy of Sciences of the United States of America, 96 (10), pp. 5575–5580.

Falcon, T., Pinheiro, D.G., Ferreira-Caliman, M.J., Turatti, I.C.C., Abreu, F.C.P., Galaschi-Teixeira, J.S., Martins, J.R., Elias-Neto, M., Soares, M.P.M., Laure, M.B., Figueiredo, V.L.C., Lopes, N.P., Simoes, Z.L.P., Garofalo, C.A. and Bitondi, M.M.G. 2019. Exploring integument transcriptomes, cuticle ultrastructure, and cuticular hydrocarbons profiles in eusocial and solitary bee species displaying heterochronic adult cuticle maturation. PloS One, 14 (3) pp. e0213796.

Farnsworth, D.R., Saunders, L.M. and Miller, A.C. 2020. A single-cell transcriptome atlas for zebrafish development. Developmental Biology, 459 (2), pp. 100–108.

Freeman, M.R. 2015. Drosophila central nervous system glia. Cold Spring Harbor Perspectives in Biology, 7 (11), pp. 10.1101/cshperspect.a020552.

Freeman, T.C., Horsewell, S., Patir, A., Harling-Lee, J., Regan, T., Shih, B.B., Prendergast, J., Hume, D.A. and Angus, T. 2022. Graphia: A platform for the graph-based visualisation and analysis of high dimensional data. PLoS Comput Biol 18(7): e1010310.

Freeman, T.C., Ivens, A., Baillie, J.K., Beraldi, D., Barnett, M.W., Dorward, D., Downing, A., Fairbairn, L., Kapetanovic, R., Raza, S., Tomoiu, A., Alberio, R., Wu, C., Su, A.I., Summers, K. M., Tuggle, C.K., Archibald, A.L. and Hume, D.A. 2012. A gene expression atlas of the domestic pig. BMC Biology, 10 pp. 90–90.

Gallai, N., Salles, J., Settele, J. and Vaissière, B.,E. 2009. Economic valuation of the vulnerability of world agriculture confronted with pollinator decline. Ecological Economics, 68 (3), pp. 810–821.

Goblirsch, M.J., Spivak, M.S. and Kurtti, T.J. 2013. A cell line resource derived from honey bee (apis mellifera) embryonic tissues. PloS One, 8 (7) pp. e69831.

Guidugli-Lazzarini, K.R., do Nascimento, A.M., Tanaka, E.D., Piulachs, M.D., Hartfelder, K., Bitondi, M.G. and Simoes, Z.L. 2008. Expression analysis of putative vitellogenin and lipophorin receptors in honey bee (apis mellifera L.) queens and workers. Journal of Insect Physiology, 54 (7), pp. 1138–1147.

Gunage, R.D., Dhanyasi, N., Reichert, H. and VijayRaghavan, K. 2017. Drosophila adult muscle development and regeneration. Seminars in Cell & Developmental Biology, 72 pp. 56–66.

He, X.J., Jiang, W.J., Zhou, M., Barron, A.B. and Zeng, Z.J. 2019. A comparison of honeybee (apis mellifera) queen, worker and drone larvae by RNA-seq. Insect Science, 26 (3), pp. 499–509.

Hepburn, H.R., Bernard, R., Davidson, B.C., Muller, W.J., Lloyd, P., Kurstjens, S.P. and Vincent, S.L. 1991. Synthesis and secretion of beeswax in honeybees. Apidologie, 22 (1), pp. 21–36.

Herranz, R., Mateos, J., Mas, J.A., Garcia-Zaragoza, E., Cervera, M. and Marco, R. 2005. The coevolution of insect muscle TpnT and TpnI gene isoforms. Molecular Biology and Evolution, 22 (11), pp. 2231–2242.

Hicks, S.C., Townes, F.W., Teng, M. and Irizarry, R.A. 2018. Missing data and technical variability in single-cell RNA-sequencing experiments. Biostatistics (Oxford, England), 19 (4), pp. 562–578.

Honeybee Genome Sequencing Consortium. 2006. Insights into social insects from the genome of the honeybee apis mellifera. Nature, 443 (7114), pp. 931–949.

Hung, R.J., Hu, Y., Kirchner, R., Liu, Y., Xu, C., Comjean, A., Tattikota, S.G., Li, F., Song, W., Ho Sui, S. and Perrimon, N. 2020. A cell atlas of the adult drosophila midgut. Proceedings of the National Academy of Sciences of the United States of America, 117 (3), pp. 1514–1523.

Jeon, J.H., Moon, K., Kim, Y. and Kim, Y.H. 2020. Reference gene selection for qRT-PCR analysis of season- and tissue-specific gene expression profiles in the honey bee apis mellifera. Scientific Reports, 10 (1), pp. 13935–4.

Kannan, K., Shook, M., Li, Y., Robinson, G.E. and Ma, J. 2019. Comparative analysis of brain and fat body gene splicing patterns in the honey bee, apis mellifera. G3 (Bethesda, Md.), 9 (4), pp. 1055–1063.

Karaiskos, N., Wahle, P., Alles, J., Boltengagen, A., Ayoub, S., Kipar, C., Kocks, C., Rajewsky, N. and Zinzen, R.P. 2017. The drosophila embryo at single-cell transcriptome resolution. Science (New York, N.Y.), 358 (6360), pp. 194–199.

Keil, T.A. 2012. Sensory cilia in arthropods. Arthropod Structure & Development, 41 (6), pp. 515–534.

King-Jones, K. and Thummel, C.S. 2005. Nuclear receptors--a perspective from drosophila. Nature Reviews.Genetics, 6 (4), pp. 311–323.

Kleijn, D., Winfree, R., Bartomeus, D., Carvalheiro, L.G., Bommarco, R., Scheper, J., Tscharntke, T., Verhulst, J. and Potts, S.G. 2015. Delivery of crop pollination services is an insufficient argument for wild pollinator conservation. Nature Communications, 6 (1), pp. 7414.

Klein, A.M., Vaissiere, B.E., Cane, J.H., Steffan-Dewenter, I., Cunningham, S.A., Kremen, C. and Tscharntke, T. 2007. Importance of pollinators in changing landscapes for world crops. Proceedings.Biological Sciences, 274 (1608), pp. 303–313.

Kubota, M., Tsuji, M., Nishimoto, M., Wongchawalit, J., Okuyama, M., Mori, H., Matsui, H., Surarit, R., Svasti, J., Kimura, A. and Chiba, S. 2004. Localization of alpha-glucosidases I, II, and III in organs of european honeybees, apis mellifera L., and the origin of alpha-glucosidase in honey. Bioscience, Biotechnology, and Biochemistry, 68 (11), pp. 2346–2352.

Leader, D.P., Krause, S.A., Pandit, A., Davies, S.A. and Dow, J.A.T. 2018. FlyAtlas 2: A new version of the drosophila melanogaster expression atlas with RNA-seq, miRNA-seq and sex-specific data. Nucleic Acids Research, 46 (D1), pp. D809–D815.

Lebestky, T., Jung, S.H. and Banerjee, U. 2003. A serrate-expressing signaling center controls drosophila hematopoiesis. Genes & Development, 17 (3), pp. 348–353.

Li, W., Evans, J.D., Huang, Q., Rodriguez-Garcia, C., Liu, J., Hamilton, M., Grozinger, C.M., Webster, T.C., Su, S. and Chen, Y.P. 2016. Silencing the honey bee (apis mellifera) naked cuticle gene (nkd) improves host immune function and reduces nosema ceranae infections. Applied and Environmental Microbiology, 82 (22), pp. 6779–6787.

Locke, M. and Krishnan, N. 1971. The distribution of phenoloxidases and polyphenols during cuticle formation. Tissue & Cell, 3 (1), pp. 103–126.

Ma, R., Rangel, J. and Grozinger, C.M. 2019. Honey bee (apis mellifera) larval pheromones may regulate gene expression related to foraging task specialization. BMC Genomics, 20 (1), pp. 592–7.

Magkrioti, C.K., Spyropoulos, I.C., Iconomidou, V.A., Willis, J.H. and Hamodrakas, S.J. 2004. cuticleDB: A relational database of arthropod cuticular proteins. BMC Bioinformatics, 5 pp. 138–138.

Menzel, R. 2012. The honeybee as a model for understanding the basis of cognition. Nature Reviews.Neuroscience, 13 (11), pp. 758–768.

Misof, B., Liu, S., Meusemann, K., Peters, R.S., Donath, A., Mayer, C., Frandsen, P.B., Ware, J., Flouri, T., Beutel, R.G., Niehuis, O., Petersen, M., Izquierdo-Carrasco, F., Wappler, T., Rust, J., Aberer, A.J., Aspock, U., Aspock, H., Bartel, D., Blanke, A., Berger, S., Bohm, A., Buckley, T.R., Calcott, B., Chen, J., Friedrich, F., Fukui, M., Fujita, M., Greve, C., Grobe, P., Gu, S., Huang, Y., Jermiin, L.S., Kawahara, A.Y., Krogmann, L., Kubiak, M., Lanfear, R., Letsch, H., Li, Y., Li, Z., Li, J., Lu, H., Machida, R., Mashimo, Y., Kapli, P., McKenna, D.D., Meng, G., Nakagaki, Y., Navarrete-Heredia, J.L., Ott, M., Ou, Y., Pass, G., Podsiadlowski, L., Pohl, H., von Reumont, B.M., Schutte, K., Sekiya, K., Shimizu, S., Slipinski, A., Stamatakis, A., Song, W., Su, X., Szucsich, N.U., Tan, M., Tan, X., Tang, M., Tang, J., Timelthaler, G., Tomizuka, S., Trautwein, M., Tong, X., Uchifune, T., Walzl, M.G., Wiegmann, B.M., Wilbrandt, J., Wipfler, B., Wong, T.K., Wu, Q., Wu, G., Xie, Y., Yang, S., Yang, Q., Yeates, D.K., Yoshizawa, K., Zhang, Q., Zhang, R., Zhang, W., Zhang, Y., Zhao, J., Zhou, C., Zhou, L., Ziesmann, T., Zou, S., Li, Y., Xu, X., Zhang, Y., Yang, H., Wang, J., Wang, J., Kjer, K.M. and Zhou, X. 2014. Phylogenomics resolves the timing and pattern of insect evolution. Science (New York, N.Y.), 346 (6210), pp. 763–767.

Morfin, N., Goodwin, P.H., Hunt, G.J. and Guzman-Novoa, E. 2019. Effects of sublethal doses of clothianidin and/or V. destructor on honey bee (apis mellifera) self-grooming behavior and associated gene expression. Scientific Reports, 9 (1), pp. 5196–0.

Nakagawa, Y. and Henrich, V.C. 2009. Arthropod nuclear receptors and their role in molting. The FEBS Journal, 276 (21), pp. 6128–6157.

Navajas, M., Migeon, A., Alaux, C., Martin-Magniette, M., Robinson, G., Evans, J., Cros-Arteil, S., Crauser, D. and Le Conte, Y. 2008. Differential gene expression of the honey bee apis mellifera associated with varroa destructor infection. BMC Genomics, 9 pp. 301–301.

Ni, J.D., Baik, L.S., Holmes, T.C. and Montell, C. 2017. A rhodopsin in the brain functions in circadian photoentrainment in drosophila. Nature, 545 (7654), pp. 340–344.

Nilsen, K.A., Ihle, K.E., Frederick, K., Fondrk, M.K., Smedal, B., Hartfelder, K. and Amdam, G.V. 2011. Insulin-like peptide genes in honey bee fat body respond differently to manipulation of social behavioral physiology. The Journal of Experimental Biology, 214 (Pt 9), pp. 1488–1497.

Nirmal, A.J., Regan, T., Shih, B.B., Hume, D.A., Sims, A.H. and Freeman, T.C. 2018. Immune cell gene signatures for profiling the microenvironment of solid tumors. Cancer Immunology Research, 6 (11), pp. 1388–1400.

Oertal, E. 1930. Metamorphosis in the honeybee. Journal of Morphology, 50 (2), pp. 295–339.

Ollerton, J., Winfree, R. and Tarrant, S. 2011. How many flowering plants are pollinated by animals? Oikos, 120 (3), pp. 321–326.

Packer, J.S., Zhu, Q., Huynh, C., Sivaramakrishnan, P., Preston, E., Dueck, H., Stefanik, D., Tan, K., Trapnell, C., Kim, J., Waterston, R.H. and Murray, J.I. 2019. A lineage-resolved molecular atlas of C. elegans embryogenesis at single-cell resolution. Science (New York, N.Y.), 365 (6459), pp. 10.1126/science.aax1971. Epub 2019 Sep 5.

Patir, A., Fraser, A.M., Barnett, M.W., McTeir, L., Rainger, J., Davey, M.G. and Freeman, T.C. 2020. The transcriptional signature associated with human motile cilia. Scientific Reports, 10 (1), pp. 10814–4.

Patir, A., Shih, B., McColl, B.W. and Freeman, T.C. 2019. A core transcriptional signature of human microglia: Derivation and utility in describing region-dependent alterations associated with alzheimer’s disease. Glia, 67 (7), pp. 1240–1253.

Raj, B., Wagner, D.E., McKenna, A., Pandey, S., Klein, A.M., Shendure, J., Gagnon, J.A. and Schier, A.F. 2018. Simultaneous single-cell profiling of lineages and cell types in the vertebrate brain. Nature Biotechnology, 36 (5), pp. 442–450.

Richardson, R.T., Ballinger, M., N, Qian, F., Christman, J.W. and Johnson, R.M. 2018. Morphological and functional characterization of honey bee, apis mellifera, hemocyte cell communitues. Apidologie, 49 pp. 397–410.

Rodriguez-Zas, S.L., Southey, B.R., Shemesh, Y., Rubin, E.B., Cohen, M., Robinson, G.E. and Bloch, G. 2012. Microarray analysis of natural socially regulated plasticity in circadian rhythms of honey bees. Journal of Biological Rhythms, 27 (1), pp. 12–24.

Ruttner, F. 1988. Taxonomy and biogeography of honey bees. Munich: Springer.

Satija, R. and Shalek, A.K. 2014. Heterogeneity in immune responses: From populations to single cells. Trends in Immunology, 35 (5), pp. 219–229.

Sebe-Pedros, A., Saudemont, B., Chomsky, E., Plessier, F., Mailhe, M.P., Renno, J., Loe-Mie, Y., Lifshitz, A., Mukamel, Z., Schmutz, S., Novault, S., Steinmetz, P.R.H., Spitz, F., Tanay, A. and Marlow, H. 2018. Cnidarian cell type diversity and regulation revealed by whole-organism single-cell RNA-seq. Cell, 173 (6), pp. 1520–1534.e20.

Seeley, T.D. 1985. Honey bee ecology. Princeton, NJ: Princeton University Press.

Shah, A.K., Kreibich, C.D., Amdam, G.V. and Munch, D. 2018. Metabolic enzymes in glial cells of the honeybee brain and their associations with aging, starvation and food response. PloS One, 13 (6) pp. e0198322.

Simpson, S.J., Sword, G.A. and Lo, N. 2011. Polyphenism in insects. Current Biology: CB, 21 (18), pp. 738.

Sinakevitch, I.T., Daskalova, S.M. and Smith, B.H. 2017. The biogenic amine tyramine and its receptor (AmTyr1) in olfactory neuropils in the honey bee (apis mellifera) brain. Frontiers in Systems Neuroscience, 11 pp. 77.

Slater, G.P., Yocum, G.D. and Bowsher, J.H. 2020. Diet quantity influences caste determination in honeybees (apis mellifera). Proceedings.Biological Sciences, 287 (1927), pp. 20200614.

Snodgrass, R.E. 1910. The anatomy of the honey bee. USA: U.S. Government Printing Office.

Sobala, L.F. and Adler, P.N. 2016. The gene expression program for the formation of wing cuticle in drosophila. PLoS Genetics, 12 (5) pp. e1006100.

Stuart, T., Butler, A., Hoffman, P., Hafemeister, C., Papalexi, E., Mauck, W.M., Hao, Y., Stoeckius, M., Smibert, P. and Satija, R. 2019. Comprehensive integration of single-cell data. Cell (Cambridge); Cell, 177 (7), pp. 1888–1902.e21.

Su, A.I., Cooke, M.P., Ching, K.A., Hakak, Y., Walker, J.R., Wiltshire, T., Orth, A.P., Vega, R. G., Sapinoso, L.M., Moqrich, A., Patapoutian, A., Hampton, G.M., Schultz, P.G. and Hogenesch, J.B. 2002. Large-scale analysis of the human and mouse transcriptomes. Proceedings of the National Academy of Sciences of the United States of America, 99 (7), pp. 4465–4470.

Tabula Muris Consortium, Overall coordination, Logistical coordination, Organ collection and processing, Library preparation and sequencing, Computational data analysis, Cell type annotation, Writing group, Supplemental text writing group and Principal investigators. 2018. Single-cell transcriptomics of 20 mouse organs creates a tabula muris. Nature, 562 (7727), pp. 367–372.

Tautz, J. 2008. The buzz about bees biology of a superorganism. 1st ed. Berlin, Heidelberg: Springer Berlin Heidelberg.

Tettamanti, G. and Casartelli, M. 2019. Cell death during complete metamorphosis. Philosophical Transactions of the Royal Society of London.Series B, Biological Sciences, 374 (1783), pp. 20190065.

Trapnell, C., Roberts, A., Goff, L., Pertea, G., Kim, D., Kelley, D.R., Pimentel, H., Salzberg, S. L., Rinn, J.L. and Pachter, L. 2012. Differential gene and transcript expression analysis of RNA-seq experiments with TopHat and cufflinks. Nature Protocols, 7 (3), pp. 562–578.

Truman, J.W. and Riddiford, L.M. 2019. The evolution of insect metamorphosis: A developmental and endocrine view. Philosophical Transactions of the Royal Society of London.Series B, Biological Sciences, 374 (1783), pp. 20190070.

Tsuchimoto, M., Aoki, M., Takada, M., Kanou, Y., Sasagawa, H., Kitagawa, Y. and Kadowaki, T. 2004. The changes of gene expression in honeybee (apis mellifera) brains associated with ages. Zoological Science, 21 (1), pp. 23–28.

Urena, E., Manjon, C., Franch-Marro, X. and Martin, D. 2014. Transcription factor E93 specifies adult metamorphosis in hemimetabolous and holometabolous insects. Proceedings of the National Academy of Sciences of the United States of America, 111 (19), pp. 7024–7029.

Van Dongen, S. 2008. Graph clustering via a discrete uncoupling process. SIAM Journal on Matrix Analysis and Applications, 30 (1), pp. 121–141.

Velarde, R.A., Robinson, G.E. and Fahrbach, S.E. 2006. Nuclear receptors of the honey bee: Annotation and expression in the adult brain. Insect Molecular Biology, 15 (5), pp. 583–595.

Verlinden, H., Vleugels, R., Marchal, E., Badisco, L., Pfluger, H.J., Blenau, W. and Broeck, J.V. 2010. The role of octopamine in locusts and other arthropods. Journal of Insect Physiology, 56 (8), pp. 854–867.

Wallberg, A., Bunikis, I., Pettersson, O.V., Mosbech, M.B., Childers, A.K., Evans, J.D., Mikheyev, A.S., Robertson, H.M., Robinson, G.E. and Webster, M.T. 2019. A hybrid de novo genome assembly of the honeybee, apis mellifera, with chromosome-length scaffolds. BMC Genomics, 20 (1), pp. 275–0.

Wang, Y., Kocher, S.D., Linksvayer, T.A., Grozinger, C.M., Page, R.E. and Amdam, G.V. 2012. Regulation of behaviorally associated gene networks in worker honey bee ovaries. The Journal of Experimental Biology, 215 (Pt 1), pp. 124–134.

Yin, L., Wang, K., Niu, L., Zhang, H., Chen, Y., Ji, T. and Chen, G. 2018. Uncovering the changing gene expression profile of honeybee (apis mellifera) worker larvae transplanted to queen cells. Frontiers in Genetics, 9 pp. 416.

Yu, G., Wang, L., Han, Y. and Qing-Yu, H. 2012. clusterProfiler: An R package for comparing biological themes among gene clusters. Omics, 16 (5), pp. 284–287.

Zammit, P.S. 2017. Function of the myogenic regulatory factors Myf5, MyoD, myogenin and MRF4 in skeletal muscle, satellite cells and regenerative myogenesis. Seminars in Cell & Developmental Biology, 72 pp. 19–32.

Zayed, A. and Robinson, G.E. 2012. Understanding the relationship between brain gene expression and social behavior: Lessons from the honey bee. Annual Review of Genetics, 46 pp. 591–615.

Zhang, F., Zhao, Y., Chao, Y., Muir, K. and Han, Z. 2013. Cubilin and amnionless mediate protein reabsorption in drosophila nephrocytes. Journal of the American Society of Nephrology: JASN, 24 (2), pp. 209–216.

Zhang, Y., Liu, X., Zhang, W. and Han, R. 2010. Differential gene expression of the honey bees apis mellifera and A. cerana induced by varroa destructor infection. Journal of Insect Physiology, 56 (9), pp. 1207–1218.

